# Mechanisms involved in cefiderocol resistance in French *Pseudomonas aeruginosa* clinical strains

**DOI:** 10.64898/2026.04.12.718081

**Authors:** Emma Gauthier, Matthieu Pisani, Maxime Bour, Mélanie Grosjean, Patrick Plesiat, Shahriar Safari, Ruben C. Hartkoorn, Léa Souro, Emma Pretot, Katy Jeannot

**Author notes:** Corresponding author. Centre National de Référence de la résistance aux antibiotiques, Laboratoire de Bactériologie, Centre Hospitalier Universitaire, 3 boulevard Fleming, 25030 Besançon, France. The authors contributed equally (Emma Gauthier, Matthieu Pisani).

## Abstract

Cefiderocol exhibits excellent *in vitro* activity against *Pseudomonas aeruginosa*; however, resistance can emerge. We investigated the molecular mechanisms underlying cefiderocol resistance (MIC >2 mg/L) in 103 clinical strains collected from 61 hospitals (2021–2024). MICs ranged from 4 to >128 mg/L, with 39.8% of strains showing MICs >8 mg/L. Although 37.8% were classified as difficult-to-treat resistant (DTR), acquired β-lactamases were detected in 72.8% of strains, including carbapenemases (39.8%), mainly NDM-1 (29.1%), and Extended Spectrum β-Lactamases (ESBLs) (38.8%). Cloning of 11 β-lactamases into pUCP24, including the acquired cephalosporinase PAC-1 and ESBLs (VEB-1, and VEB-9), resulted in marked increases in cefiderocol MICs (up to 128-fold). Introduction of 6 mutations in the PDC enzyme into a PAO1Δ*bla*_PDC-1_ background increased MICs up to 4 mg/L and conferred cross-resistance to ceftolozane/tazobactam, notably F121L, G157D, T70I, and E219K. Alterations in siderophore transporters or regulators were identified in 38.8% of strains, most frequently a PirR frameshift (R132*fs*), consistent with PirR inactivation, which was confirmed in the PAO1 strain to contribute to cefiderocol resistance. Overall, cefiderocol resistance in clinical strains is multifactorial, mainly involving acquired β-lactamases (ESBLs, carbapenemases) and impaired siderophore uptake (PiuA/PiuD, PirA, PiuC), leading to high-level resistance (>8 mg/L). The polyclonal distribution and diversity of mechanisms highlight the need for routine susceptibility testing and surveillance. Detection of NDM producers is critical, as cefiderocol should be used with caution in this context.

## Introduction

*Pseudomonas aeruginosa* is notoriously known as the most common cause of healthcare-associated pneumonia in intensive care units (20.1% in Europe) (1). While most isolates (67.6%) remain susceptible to first-line antibiotics such as ceftazidime, piperacillin/tazobactam, and meropenem, 11.3% of invasive European isolates are Multidrug resistant (MDR), extensively resistant (XDR) or difficult to treat resistance (DTR) limiting therapeutic options to a few antibiotics including cefiderocol (2). This first cephalosporin-siderophore is currently a therapeutic option in adults to combat *P. aeruginosa* infections associated with isolates resistant to carbapenems, ceftazidime/avibactam ceftolozane/tazobactam, and imipenem/cilastatin/relebactam (3). While this cephalosporin-siderophore generally has an excellent *in vitro* activity against such *P. aeruginosa* clinical isolates, several studies have reported several intrinsic and acquired mechanism of resistance to cefiderocol (4–6) In relation to its active entry into the periplasmic space, loss of function mutation in the TonB-dependent siderophore transporters PiuA or PiuD (depending on the genetic background of the strain) have been reported to lead to 16- and 32-fold increase in cefiderocol MIC in both the PAO1 reference strain, as well in a clinical strain after a ceftazidime/avibactam exposure (7–10). Recently, mutations of the cognate TonB-receptor FptA (the receptor for pyochelin, the second endogenous *P. aeruginosa* siderophore) were associated with a 2- to 4-fold decrease in cefiderocol susceptibility in the PA14 reference strain (11). In addition, while cefiderocol is active against *P. aeruginosa* AmpC overproducing strains, certain mutations in AmpC (such as T96I, F121L + G222S, G183D, G214R, E219K, E247K, and L320P), when expressed recombinantly in PAO1 or in PAO1Δ*ampC*Δ*oprD*, have been shown to impact cefiderocol in *vitro* activity (12, 13). Of these PDC variants, the E247K mutation has also been reported in two clinical cefiderocol resistant strains of *P. aeruginosa* (with MIC values of 8 and 32 mg/L) according to EUCAST breakpoints (MIC >2 mg/L) following exposure to ceftolozane/tazobactam (14). Additionally, a 4 amino-acids deletion (TPMA) at positions 316-319 of PDC was identified in a clinical strain of *P. aeruginosa* (patient treated previously with cefiderocol), that led to a significant increase in the cefiderocol MIC (from 0.12 to 2 mg/L) (15). Beyond mutations driven intrinsic resistance mechanisms, various types of ß-lactamases, when expressed recombinantly, have been shown to mediate cefiderocol resistance, including carbapenemases (PER-7, NDM-1, and SPM-1) as well as extended-spectrum ß-lactamases (PER-1, −6; SHV-2a, −12; and BEL-1, −2) or oxacillinases derived from OXA-2 (OXA-2, −15, −161,-226, −539, −681, −737, −935), OXA-10 (OXA-14, −19, −21, −794, −795,−824, −836,) and OXA-46 (OXA-46, −779) (13, 16, 17). Despite the various potential mechanism of cefiderocol resistance identified, there have been limited reports on the diversity of such mechanisms in cefiderocol-resistant *P. aeruginosa* clinical strains (4, 18). The aim of this study was to map the genetic mechanisms contributing to cefiderocol resistance (MIC>2 mg/L) in a French collection of 103 cefiderocol resistance *P. aeruginosa* clinical strains.

## Results

From the collection of the FNRC for antibiotic resistance *in P. aeruginosa*, 103 clinical strains (covering 4-years) were considered resistant to cefiderocol according to its clinical breakpoints defined by EUCAST 2025 (MIC>2 mg/L). These strains were originally isolated from the respiratory tract (29.1%), blood (22.3%), urine (18.4%), rectal swabs (9.7%), bone (8.7%), skin (6.8%) and other sites (5.0%) (Table S1). The cefiderocol susceptibility of these strains ranged from an MIC of 4 (*n*=35, 34.0%) to ≥128 mg/L (*n*=8, 7.8%), with 41/103 strains (39.8%) having an MIC values higher than 8 mg/L, which corresponds to the cefiderocol resistance according to CLSI breakpoints 2025 (19) (Table 1). The cefiderocol-resistant strains were highly resistant to first line antibiotics (piperacillin/tazobactam 90.3%, ceftazidime 95.1%, and cefepime 96.1%) and carbapenems (imipenem 81.5%, and meropenem 71.8%) as well as to combinations of ß-lactam and ß-lactamase inhibitors (ceftolozane/tazobactam 90.3%, ceftazidime/avibactam 89.3%) (Tables 1 and S1). Aztreonam (66.0%) and imipenem/relebactam (68.9%) were the only ß-lactams for which resistance rates were found to be below 70%. When integrating fluoroquinolones susceptibility into the analysis, a total of 39 cefiderocol resistant strains (37.8%) could be classified as Difficult-to-Treat Resistant (DTR) (20). These DTR strains were also resistant to ceftolozane/tazobactam (35/39 DTR strains), ceftazidime/avibactam (30/39 DTR strains), imipenem/relebactam (36/39 DTR strains), and amikacin (33/39 DTR strains), though most remained sensitive to colistin (resistance found in 2/39 DTR strains).

**Table 1.**
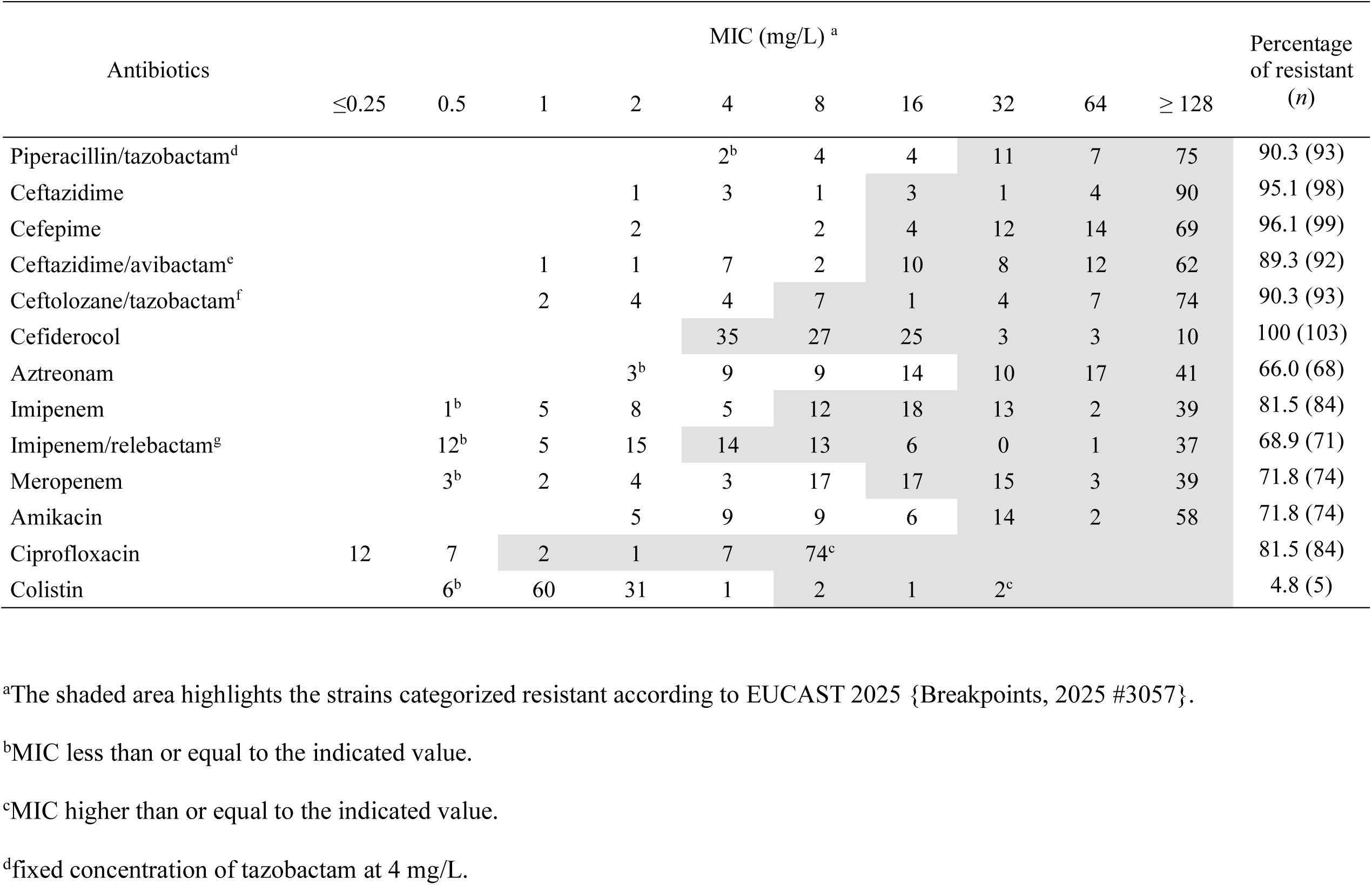
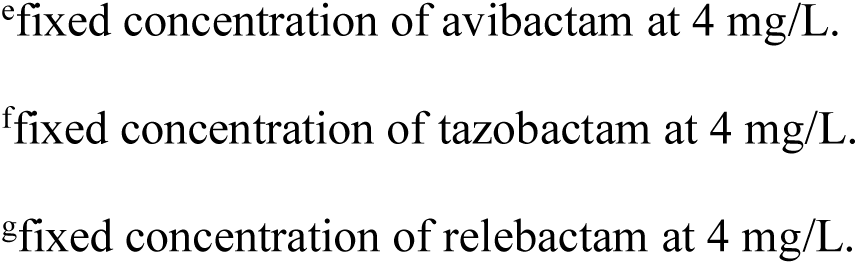
Antibiotic MIC distribution of 103 cefiderocol-resistant clinical strains of *P. aeruginosa*.

| Antibiotics | MIC (mg/L) <sup>a</sup> |  |  |  |  |  |  |  |  |  | Percentage<br>of resistant<br>(n) |
| --- | --- | --- | --- | --- | --- | --- | --- | --- | --- | --- | --- |
|  | ≤0.25 | 0.5 | 1 | 2 | 4 | 8 | 16 | 32 | 64 | ≥128 |  |
| Piperacillin/tazobactam<sup>d</sup> |  |  |  |  | 2<sup>b</sup> | 4 | 4 | 11 | 7 | 75 | 90.3 (93) |
| Ceftazidime |  |  |  | 1 | 3 | 1 | 3 | 1 | 4 | 90 | 95.1 (98) |
| Cefepime |  |  |  | 2 |  | 2 | 4 | 12 | 14 | 69 | 96.1 (99) |
| Ceftazidime/avibactam<sup>e</sup> |  |  | 1 | 1 | 7 | 2 | 10 | 8 | 12 | 62 | 89.3 (92) |
| Ceftolozane/tazobactam<sup>f</sup> |  |  | 2 | 4 | 4 | 7 | 1 | 4 | 7 | 74 | 90.3 (93) |
| Ceftiderocol |  |  |  |  | 35 | 27 | 25 | 3 | 3 | 10 | 100 (103) |
| Aztreonam |  |  |  | 3<sup>b</sup> | 9 | 9 | 14 | 10 | 17 | 41 | 66.0 (68) |
| Imipenem |  | 1<sup>b</sup> | 5 | 8 | 5 | 12 | 18 | 13 | 2 | 39 | 81.5 (84) |
| Imipenem/relebactam<sup>g</sup> |  | 12<sup>b</sup> | 5 | 15 | 14 | 13 | 6 | 0 | 1 | 37 | 68.9 (71) |
| Meropenem |  | 3<sup>b</sup> | 2 | 4 | 3 | 17 | 17 | 15 | 3 | 39 | 71.8 (74) |
| Amikacin |  |  |  | 5 | 9 | 9 | 6 | 14 | 2 | 58 | 71.8 (74) |
| Ciprofloxacin | 12 | 7 | 2 | 1 | 7 | 74<sup>c</sup> |  |  |  |  | 81.5 (84) |
| Colistin |  | 6<sup>b</sup> | 60 | 31 | 1 | 2 | 1 | 2<sup>c</sup> |  |  | 4.8 (5) |

<sup>a</sup>The shaded area highlights the strains categorized resistant according to EUCAST 2025 {Breakpoints, 2025 #3057}.
<sup>b</sup>MIC less than or equal to the indicated value.
<sup>c</sup>MIC higher than or equal to the indicated value.
<sup>d</sup>fixed concentration of tazobactam at 4 mg/L.
<sup>e</sup>fixed concentration of avibactam at 4 mg/L.
<sup>f</sup>fixed concentration of tazobactam at 4 mg/L.
<sup>g</sup>fixed concentration of relebactam at 4 mg/L.

**Table 2.**
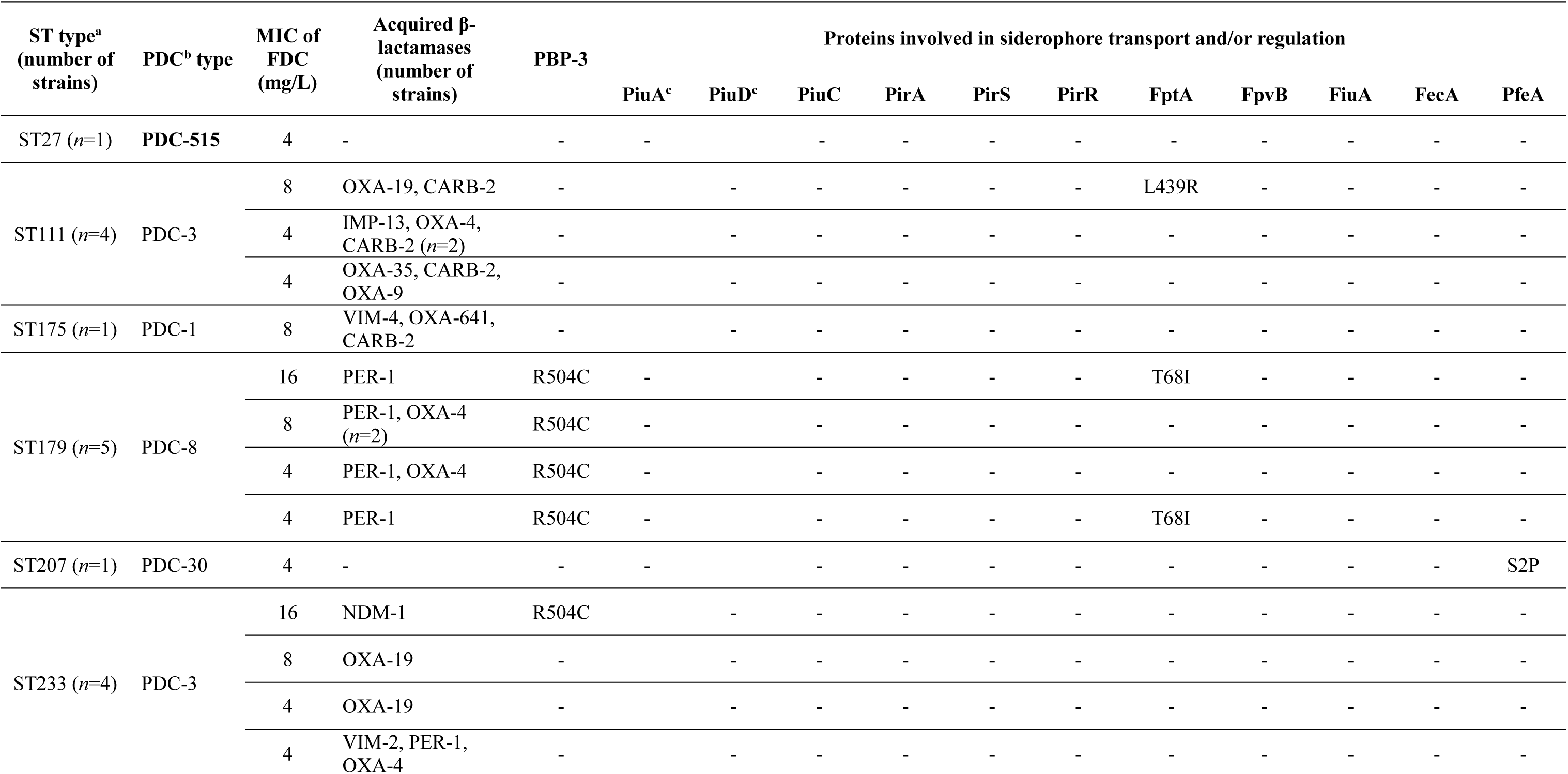
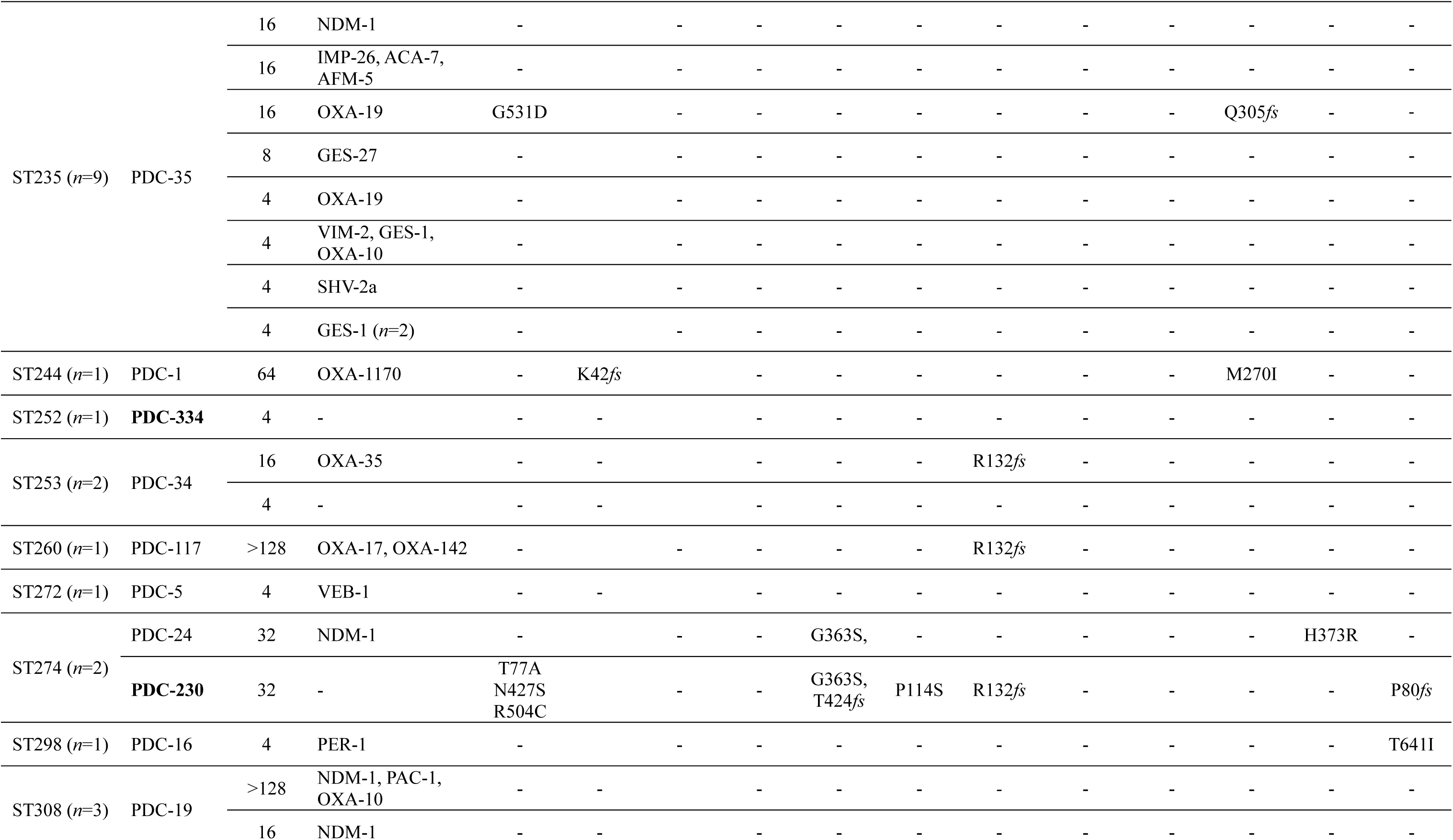
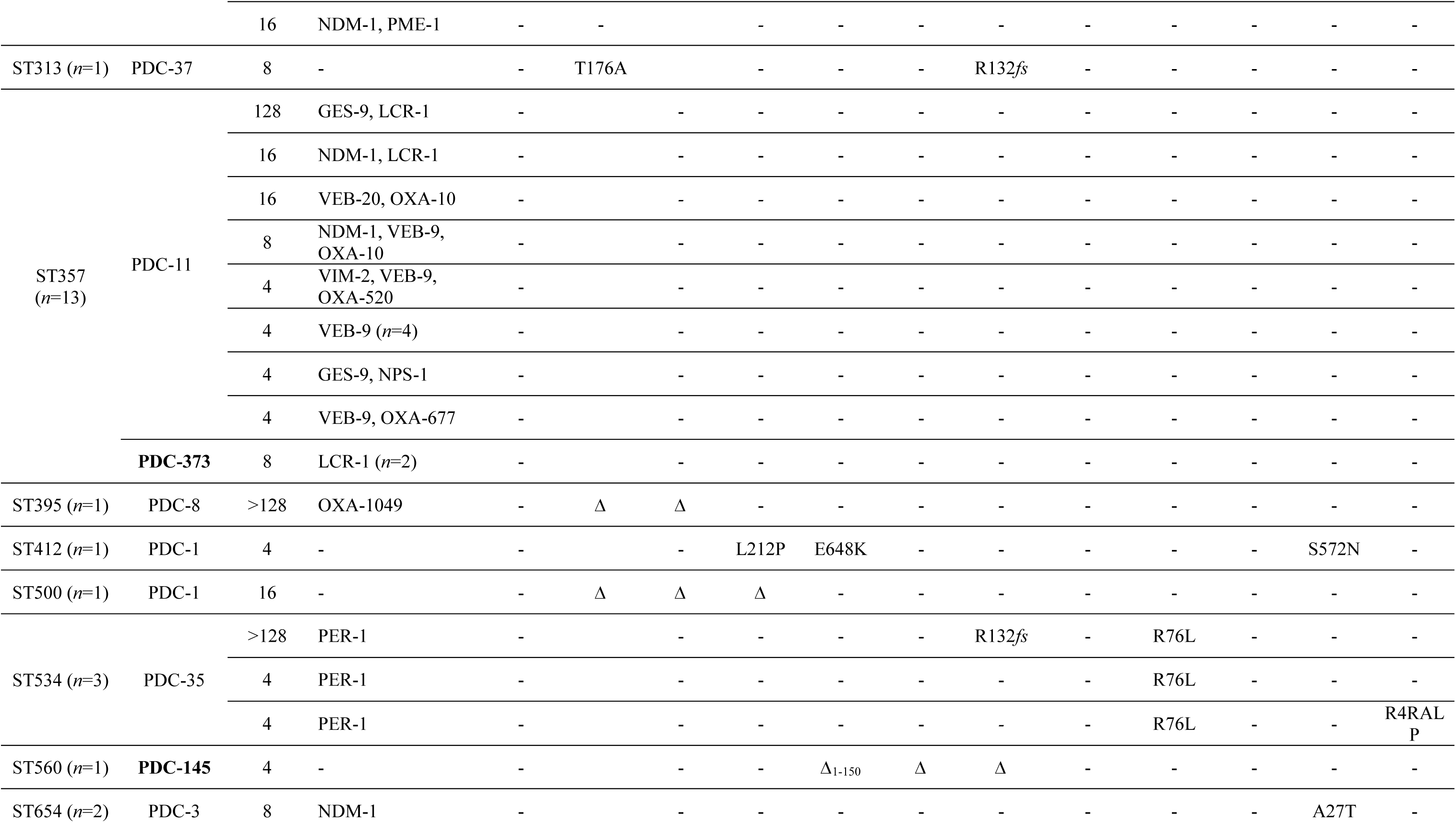
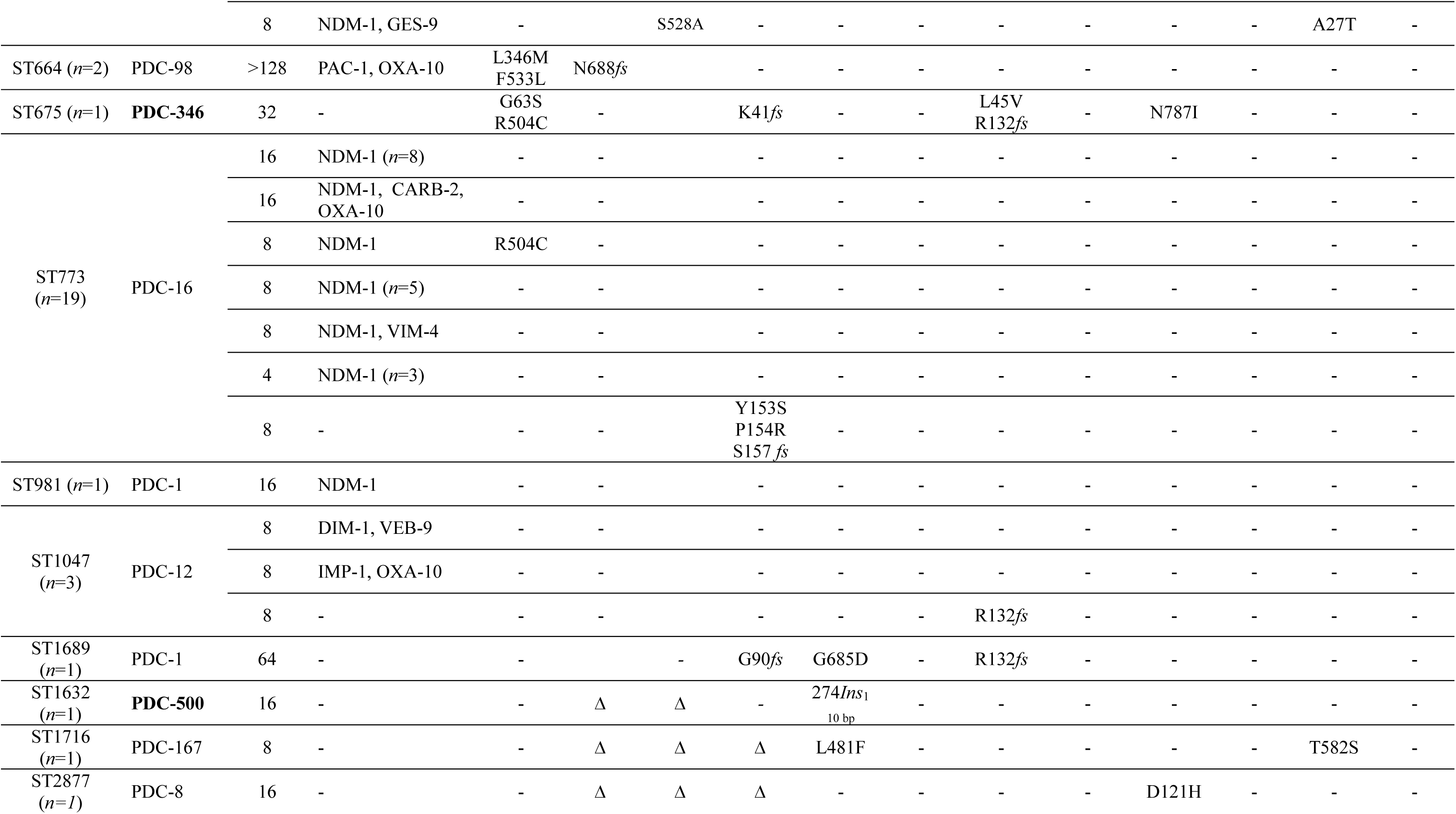
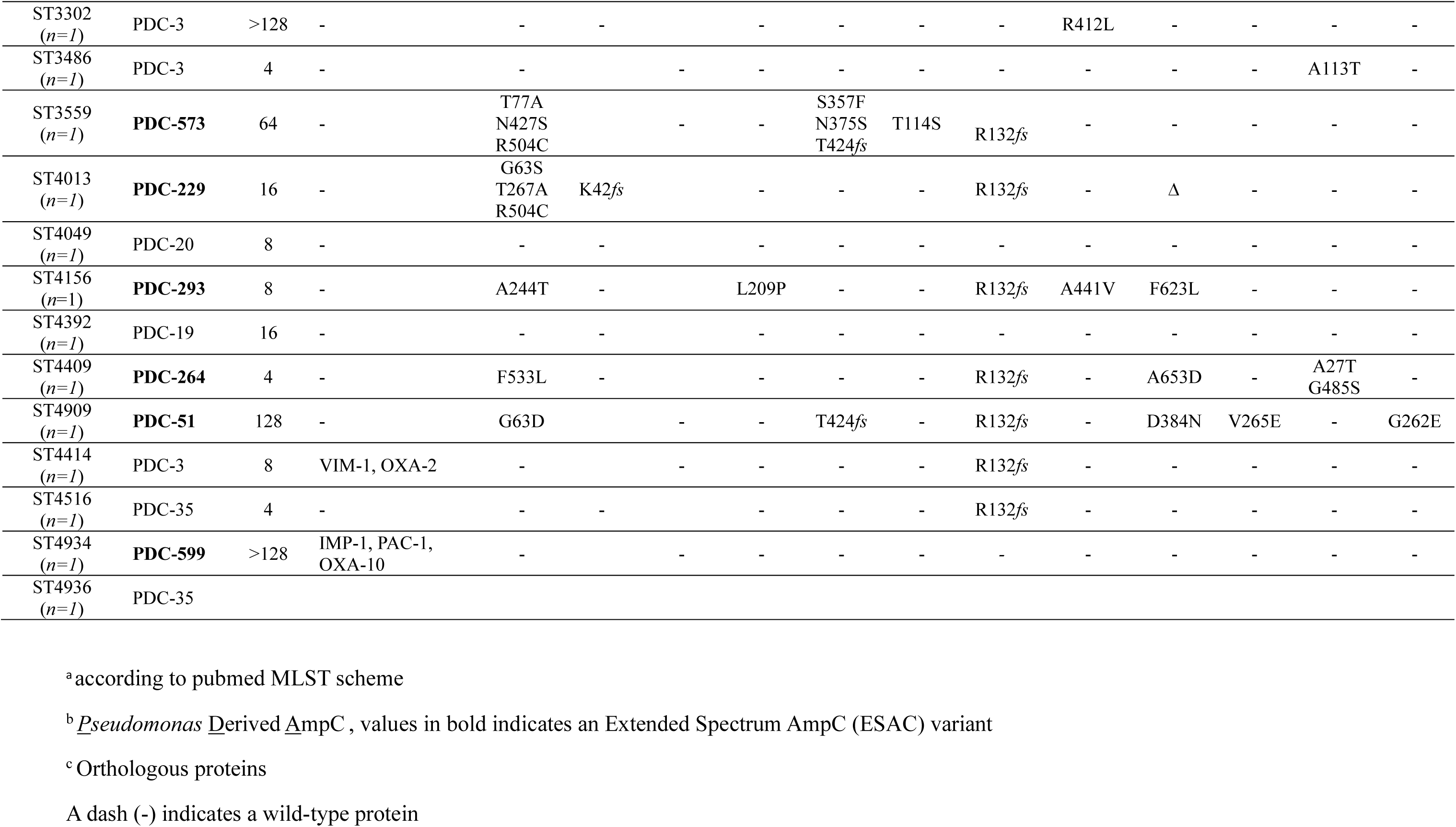
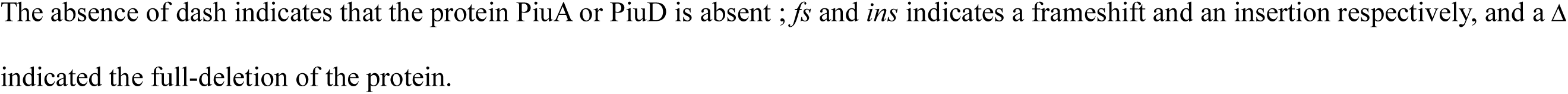
Resistance mechanisms identified in cefiderocol resistant (MIC >2 mg/L) *Pseudomonas aeruginosa* clinical strains (*n*=103).

When examining the Sequence Type (ST) of these strains, a total of 45 STs were identified, with only 6 STs represented by more than 3 strains [specifically: ST111 (*n*=4), ST179 (*n*=5), ST233 (*n*=4), ST235 (*n*=8), ST357 (*n*=13), and ST773 (*n*=19)] (Fig 1). With the exception of ST179 and ST773, the STs associated with cefiderocol-resistant strains were previously classified among the top 10 high-risk lineages found to accumulate resistance determinants (21). Overall, the cefiderocol-resistant strains were polyclonal, with cefiderocol susceptibility not correlating with ST, which infers that the probability of cefiderocol resistance cannot be predicted in routine clinical laboratories by ST determination.

**Fig 1.**
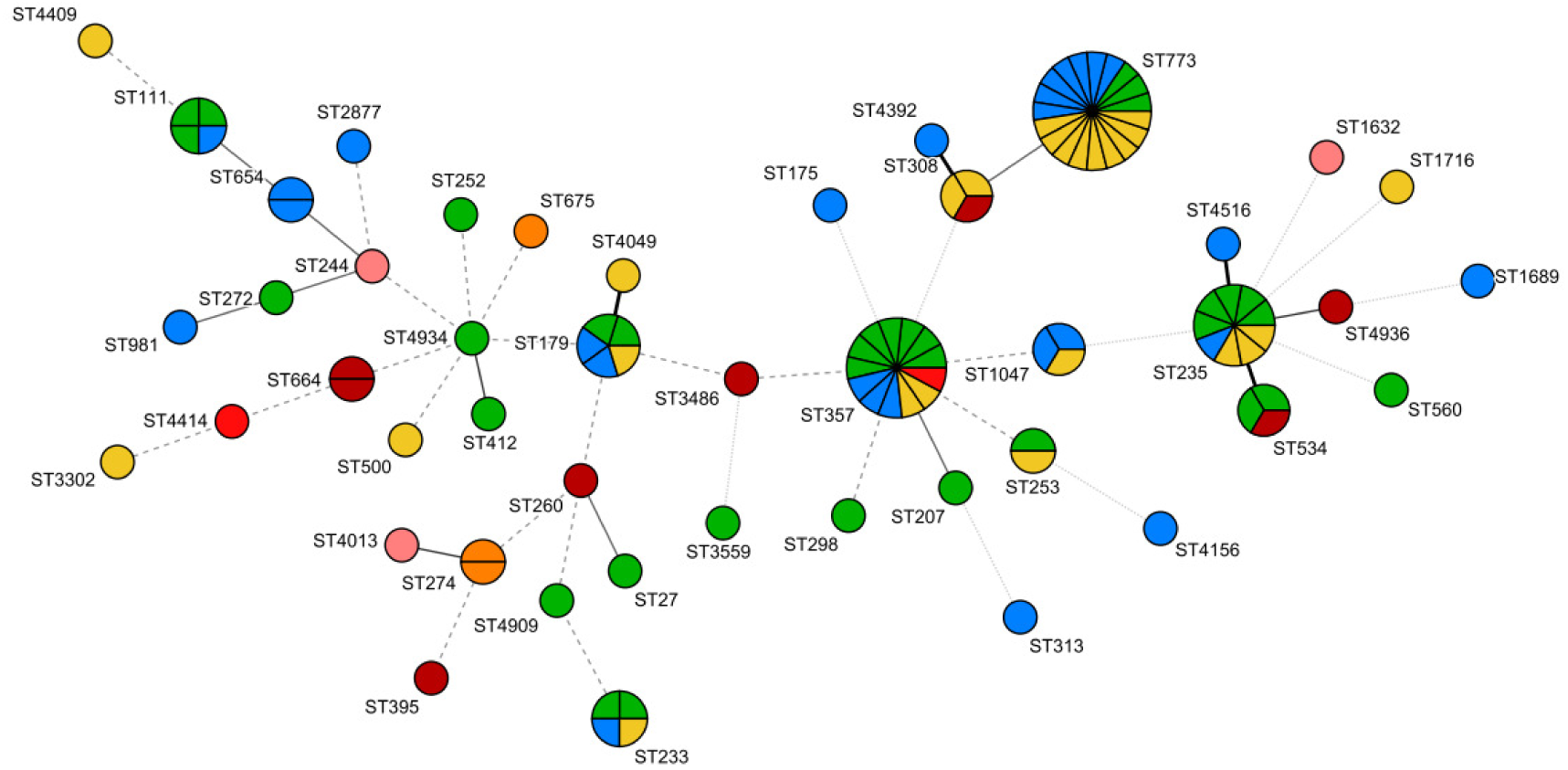
Sequence Type (ST) distribution of 103-cefiderocol-resistant strains. The MIC values of cefiderocol for the studied strains are represented as followed: green for 4 μg/mL, blue for 8 μg/mL, yellow for 16 μg/mL, orange for 32 μg/mL, pink for 64 μg/mL, red for ≥ 128 μg/mL. The lines connecting the STs represent the number of allelic differences between them. A solid line, a stippled line, and a very fine dotted line, indicate 2, 3 and 4 allelic differences between STs, respectively. The minimum spanning tree was drawn on BioNumerics 7.1 and modified on Inkscape 1.4.2 softwares.

### Transferable resistance mechanisms in cefiderocol-resistant clinical strains

Whole-genome sequencing data was used to map the acquisition of transferable β-lactamases in the 103 cefiderocol resistant strains. Of these strains a total of 77/103 (74.7%) were found to have acquired at least one transferable ß-lactamase, including carbapenemases (41/103 strains, 39.8%), ESBL (40/103 strains, 38.8%), penicillinases (23/103 strains, 22.3%) and cephalosporinase PAC-1 (4/103, 3.8%) (Tables 2 and S2). Twenty-nine cefiderocol resistant strains (28.1%) were identified to carry multiple ß-lactamases, of with 12 strains carried three such degrading enzymes (9.7%). Among these, 9/29 strains carried a carbapenemase and an ESBL, and 3/29 strains carried two distinct carbapenemases (AFM-5/IMP-26, VIM-4/OXA-641, and VIM-4/NDM-1). Of the 41 carbapenemase-producing strains, metallo-β-lactamases were the most commonly identified, specifically NDM-1 (*n*=30), IMP-1 (*n*=2), IMP-13 (*n*=2), IMP-26 (*n*=1), VIM-1 (*n*=1), VIM-2 (*n*=3), VIM-4 (*n*=2), AFM-2 (*n*=1), and DIM-1 (*n*=1), with only one strains identified to carry a oxacillinase (OXA-641) with carbapenemase activity.

While nearly one-third (29.1%, *n*=30) of cefiderocol-resistant strains produced NDM-1, their cefiderocol susceptibility ranged from an MIC of 4 to >128 mg/L, indicating that other factors may be influencing susceptibility. These factors could include the expression of additional ß-lactamases, with seven NDM-1 producer strains also found to carry either one (*n*=4) or two additional ß-lactamases (*n*=3), beit, PAC-1, OXA-10, LCR-1, CARB-2, PME-1, VEB-9, or GES-9 (Tables 2 and S2). Of the many MBL identified in the cefiderocol resistant strains, the majority have previously been validated to cause cefiderocol resistance in engineered laboratory strains *in vitro* (22–24). An exception to this are four MBL-enzymes (IMP-1, IMP-26, VIM-4, and AFM-5) whose role in cefiderocol resistance has not been previously confirmed (22–24), whose role in cefiderocol resistance has not been previously confirmed. To evaluate if these MBLs are involved in conferring cefiderocol resistance, a broad-spectrum, zinc chelating MBL inhibitor, dipicolinic acid, was used to inactivate these MBLs in the cefiderocol resistant strains, and evaluate cefiderocol susceptibility (Table 3).

With the exception of the clinical strain producing DIM-1, for which no effect was expected based on previous *in vitro* observations, the addition of dipicolinic acid fully or partially restored cefiderocol susceptibility in these MBL (NDM-1/VIM-4, VIM-2/GES-1/OXA-10) producers (Table 3). These data support the role of VIM-2/GES-1/OXA-10 in cefiderocol resistance in clinical strains of *P. aeruginosa* (Table 3).

Amongst the 103 cefiderocol resistant strains there was a high proportion of clinical strains carrying ESBLs genes (38.8%, *n*=40), which represents a higher-than-average prevalence in clinical strains in France (25). Amongst the ESBL, Ambler Class A ESBLs including were the most frequent, including SHV-2a (*n*=1), PER-1 (*n*=10), PME-1 (*n*=1), VEB-1 (*n*=1), VEB-9 (*n*=8), VEB-20 (*n*=1), GES-1 (*n*=3), GES-9 (*n*=3), GES-27 (*n*=1), and ACA-7 (*n*=1). These were followed by Ambler Class D ES-OXA enzymes (*n*=12), among which OXA-19 was the most common variant (*n*=5) (Tables 2 and S2). While the role of some of ESBL has been confirmed in cefiderocol resistance, this is not the case for all. To confirm their contribution to cefiderocol resistance VEB-1, VEB-9, ACA-7, OXA-142, OXA-1049, and OXA-1170 (and others) were cloned into a pUCP24 plasmid and expressed in PAO1, followed by antibiotic susceptibility testing (Table 4). Data confirmed all these ESBL enzymes to confer 4- to 128-fold cefiderocol resistance with particularly high resistance mediated by VEB-9 and VEB-1 (Table 4).

**Table 3.** Impact of dipicolinic acid on cefiderocol susceptibility in MBL-producing *P. aeruginosa* clinical strains.

| Clinical Strain | Acquired β-lactamase | MIC of | |
| --- | --- | --- | --- |
|  |  | cefiderocol (mg/L) | cefiderocol (mg/L) / dipicolinic acid (100 mg/L) |
| CIP76110 | Reference strain | 0.125 | 0.125 |
| PSA-19 | **NDM-1** | 32 | 0.25 |
| 9159 | **NDM-1/VIM-4** | 8 | 0.06 |
| 9611 | **VIM-4/OXA-641/CARB-2** | 8 | 1 |
| 11334 | **IMP-1/OXA-10** | 8 | 2 |
| 11558 | **IMP-1/PAC-1/OXA-10** | >128 | 8 |
| 8610 | **IMP-13/CARB-2/OXA-4** | 4 | 1 |
| 9321 | **IMP-13/CARB-2/OXA-4** | 4 | 1 |
| 11485 | **IMP-26/AFM-5/ACA-7** | 16 | 0.5 |
| 11016 | **VIM-1/OXA-2** | 8 | 2 |
| 9350 | **VIM-2/GES-1/OXA-10** | 4 | 0.125 |
| 11165 | **VIM-2/VEB-9/OXA-520** | 4 | 1 |
| 11595 | **VIM-2**/PER-1/OXA-4 | 4 | 0.25 |
| 10311 | **DIM-1**/VEB-9 | 8 | 8 |

The metallo-β-lactamase are indicated in bold.

**Table 4.**
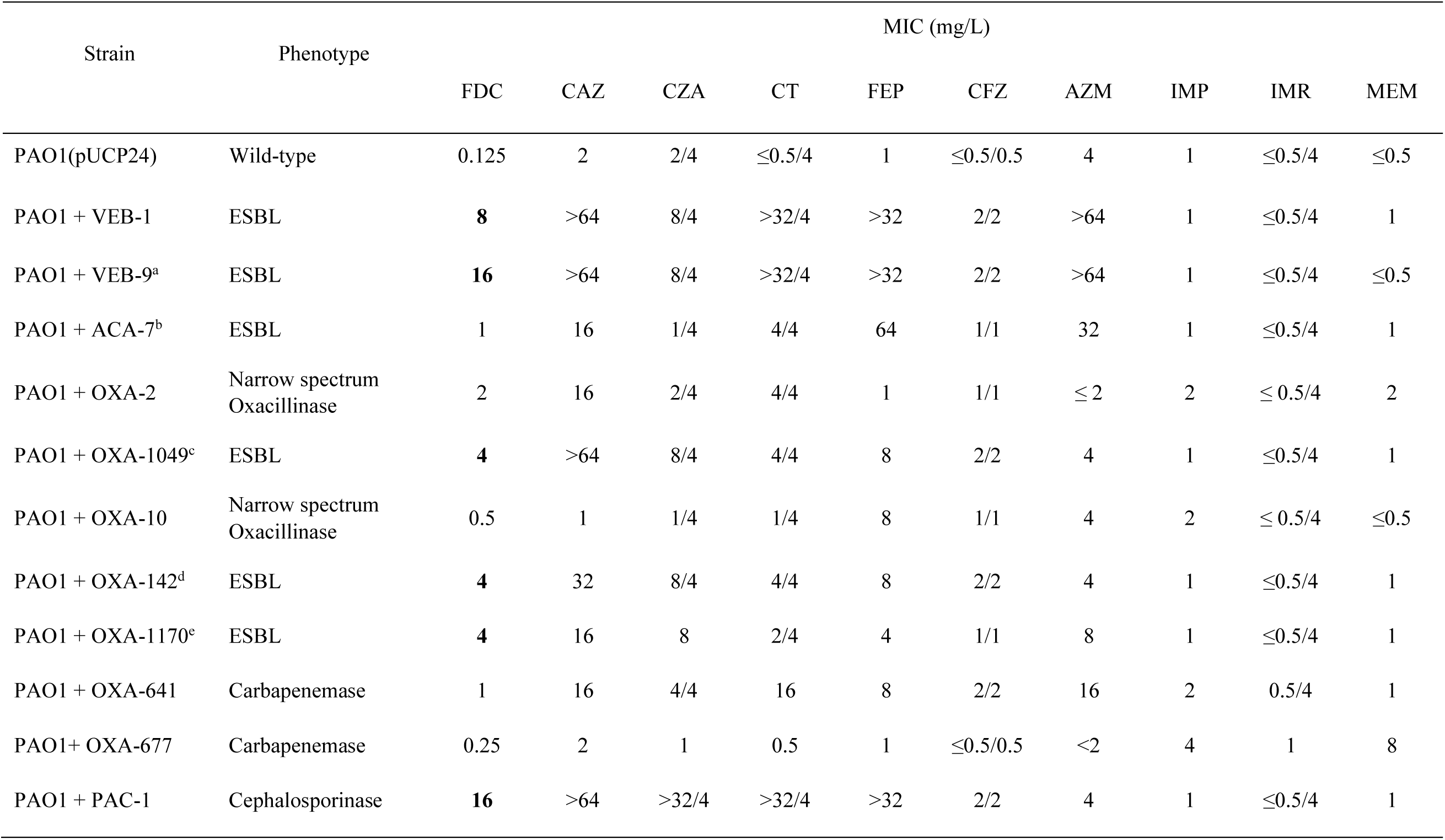
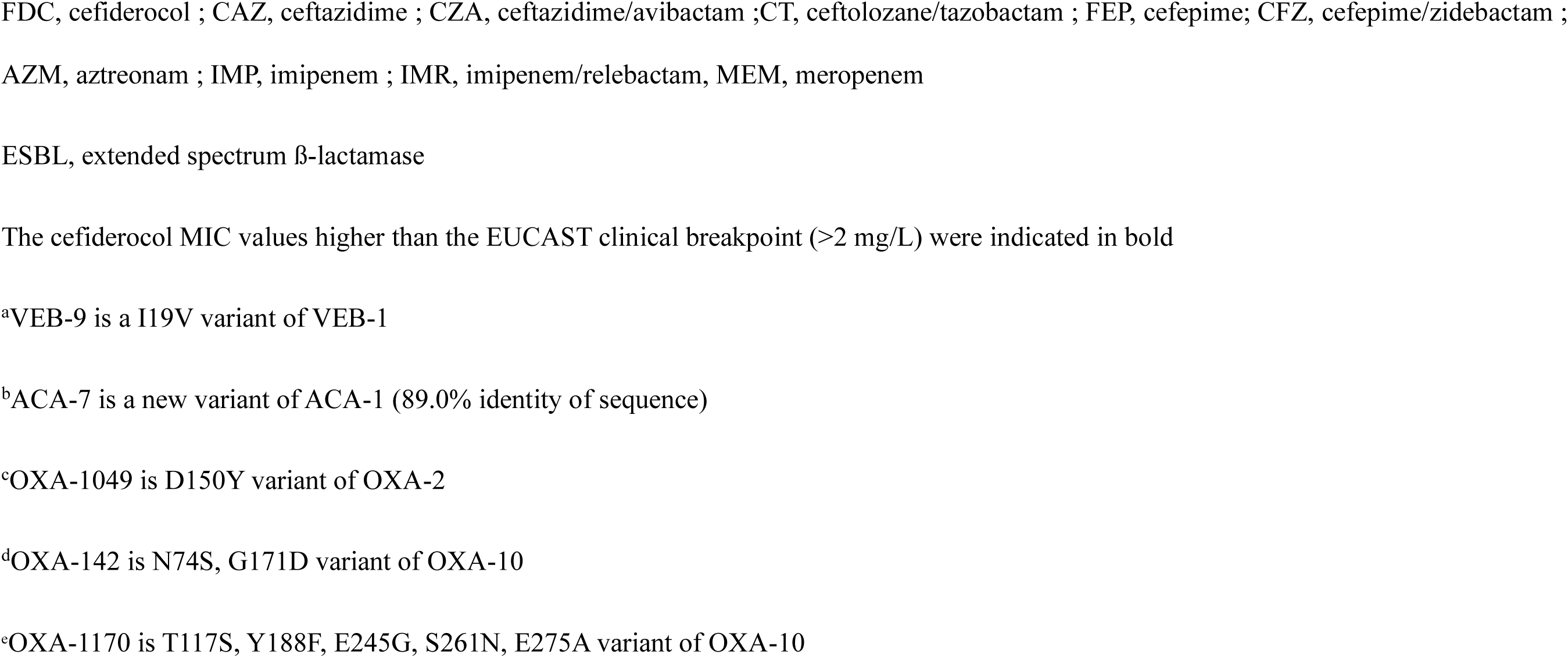
Antibiotic susceptibility of PAO1 carrying recombinant ß-lactamase plasmids.

| Strain | Phenotype | MIC (mg/L) |  |  |  |  |  |  |  |  |  |
| --- | --- | --- | --- | --- | --- | --- | --- | --- | --- | --- | --- |
|  |  | FDC | CAZ | CZA | CT | FEP | CFZ | AZM | IMP | IMR | MEM |
| PAO1(pUCP24) | Wild-type | 0.125 | 2 | 2/4 | $\leq 0.5/4$ | 1 | $\leq 0.5/0.5$ | 4 | 1 | $\leq 0.5/4$ | $\leq 0.5$ |
| PAO1 + VEB-1 | ESBL | **8** | >64 | 8/4 | >32/4 | >32 | 2/2 | >64 | 1 | $\leq 0.5/4$ | 1 |
| PAO1 + VEB-9<sup>a</sup> | ESBL | **16** | >64 | 8/4 | >32/4 | >32 | 2/2 | >64 | 1 | $\leq 0.5/4$ | $\leq 0.5$ |
| PAO1 + ACA-7<sup>b</sup> | ESBL | 1 | 16 | 1/4 | 4/4 | 64 | 1/1 | 32 | 1 | $\leq 0.5/4$ | 1 |
| PAO1 + OXA-2 | Narrow spectrum Oxacillinase | 2 | 16 | 2/4 | 4/4 | 1 | 1/1 | $\leq 2$ | 2 | $\leq 0.5/4$ | 2 |
| PAO1 + OXA-1049<sup>c</sup> | ESBL | **4** | >64 | 8/4 | 4/4 | 8 | 2/2 | 4 | 1 | $\leq 0.5/4$ | 1 |
| PAO1 + OXA-10 | Narrow spectrum Oxacillinase | 0.5 | 1 | 1/4 | 1/4 | 8 | 1/1 | 4 | 2 | $\leq 0.5/4$ | $\leq 0.5$ |
| PAO1 + OXA-142<sup>d</sup> | ESBL | **4** | 32 | 8/4 | 4/4 | 8 | 2/2 | 4 | 1 | $\leq 0.5/4$ | 1 |
| PAO1 + OXA-1170<sup>e</sup> | ESBL | **4** | 16 | 8 | 2/4 | 4 | 1/1 | 8 | 1 | $\leq 0.5/4$ | 1 |
| PAO1 + OXA-641 | Carbapenemase | 1 | 16 | 4/4 | 16 | 8 | 2/2 | 16 | 2 | 0.5/4 | 1 |
| PAO1+ OXA-677 | Carbapenemase | 0.25 | 2 | 1 | 0.5 | 1 | $\leq 0.5/0.5$ | <2 | 4 | 1 | 8 |
| PAO1 + PAC-1 | Cephalosporinase | **16** | >64 | >32/4 | >32/4 | >32 | 2/2 | 4 | 1 | $\leq 0.5/4$ | 1 |

FDC, cefiderocol ; CAZ, ceftazidime ; CZA, ceftazidime/avibactam ; CT, ceftolozane/tazobactam ; FEP, cefepime; CFZ, cefepime/zidebactam ; AZM, aztreonam ; IMP, imipenem ; IMR, imipenem/relebactam, MEM, meropenem
ESBL, extended spectrum β-lactamase
The cefiderocol MIC values higher than the EUCAST clinical breakpoint ( $>2$ mg/L) were indicated in bold
<sup>a</sup>VEB-9 is a I19V variant of VEB-1
<sup>b</sup>ACA-7 is a new variant of ACA-1 (89.0% identity of sequence)
<sup>c</sup>OXA-1049 is D150Y variant of OXA-2
<sup>d</sup>OXA-142 is N74S, G171D variant of OXA-10
<sup>e</sup>OXA-1170 is T117S, Y188F, E245G, S261N, E275A variant of OXA-10

In addition to ESBL enzymes, four highly cefiderocol resistant clinical strains (CMI>128 mg/L) were found to carry a PAC-1 cephalosporinase in association with an OXA-10 oxacillinase or, the MBL, IMP-1 or NDM-1 carbapenemases (26). As above, to explore the specific role of PAC-1 cephalosporinase in cefiderocol resistance, *bla*_PAC-1_ was cloning into PAO1, and found to increase in cefiderocol MIC, from 0.125 to 16 mg/L (Table 4). These data reveal that carbapenemases (particularly NDM-1), as well as class A and D ESBLs, significantly contribute as determinants of cefiderocol susceptibility.

### Intrinsic enzymatic resistance mechanisms in cefiderocol-resistant clinical strains

Amongst the 103 resistant strains, 30 different PDC sequences were identified. The most frequent variants were PDC-16 (19.4%, *20/103*), PDC-35 (13.6%, *14/103*), PDC-3 (12.6%, *13/103*), PDC-11 (10.7%, *11/103*), and PDC-8 (6.8%, *7/103*). Genomic analysis of the PDCs, and exclusion of natural polymorphisms, identified 14 strains exhibited an ESAC variants (13.6%), including mutations V211A (*n*=5), E219K (*n*=3), T70I (*n*=2), G156D (*n*=2), F121L (*n*=1), and Δ291-293 (Table 5). To evaluate the contribution of these mutations, the different *bla*_PDC-1_ mutants were cloned into PAO1Δ*bla*_PDC-1_, and their impact on cefiderocol susceptibility determined. The substitution E219K and the deletion Δ292-294, located in Ω loop and the R2 domains of the proteins respectively, led to the largest increase in cefiderocol MIC (from 0.5 to 4 mg/L) in comparison with the other substitutions, which resulted in MICs of 1, and 2 mg/L. In addition to cefiderocol, most of these mutations significantly affected susceptibility to ceftolozane/tazobactam, particularly E219K, and T70I (Table 5), indicating a cross-resistance between ceftolozane/tazobactam and cefiderocol.

**Table 5.**
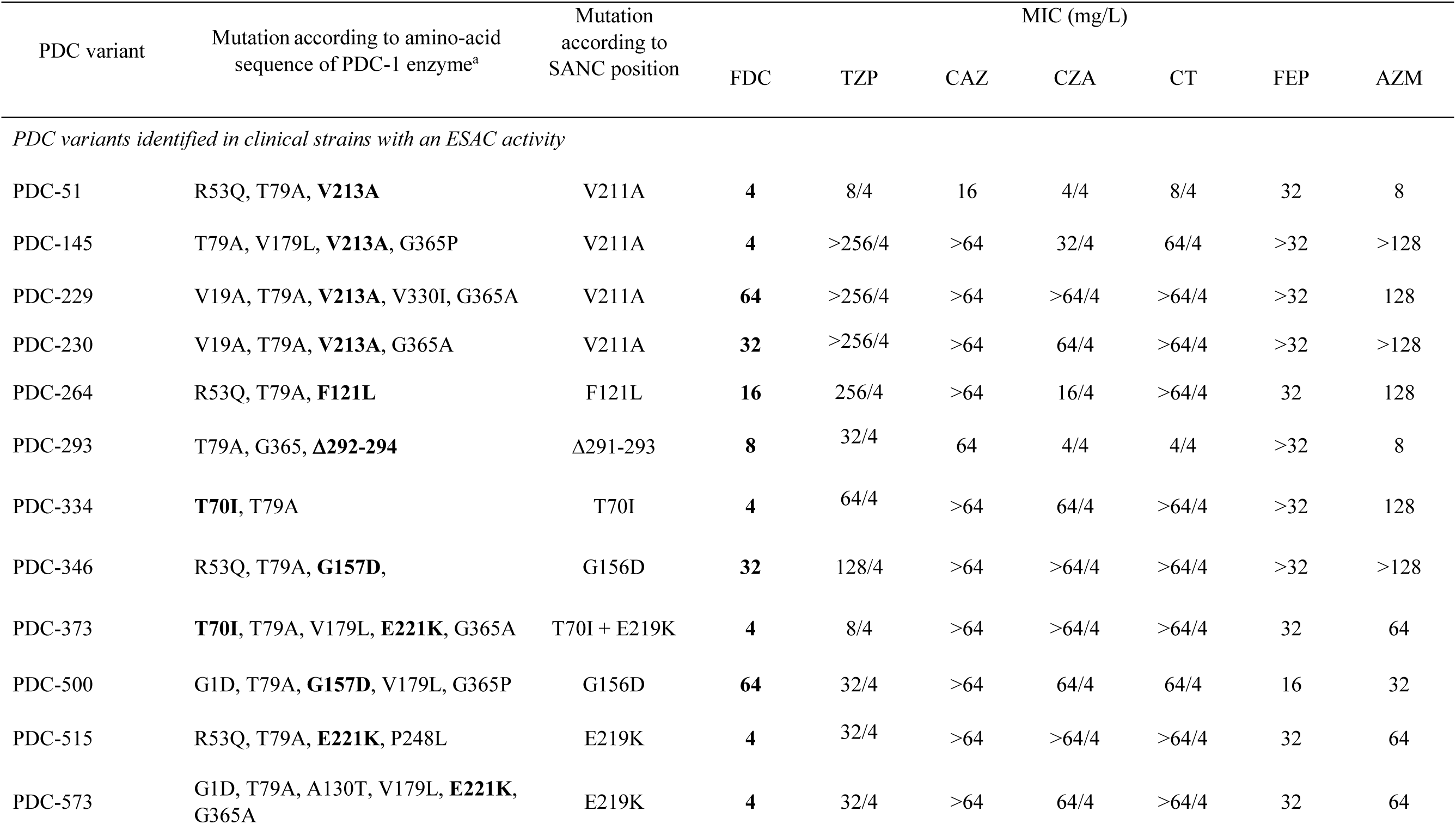
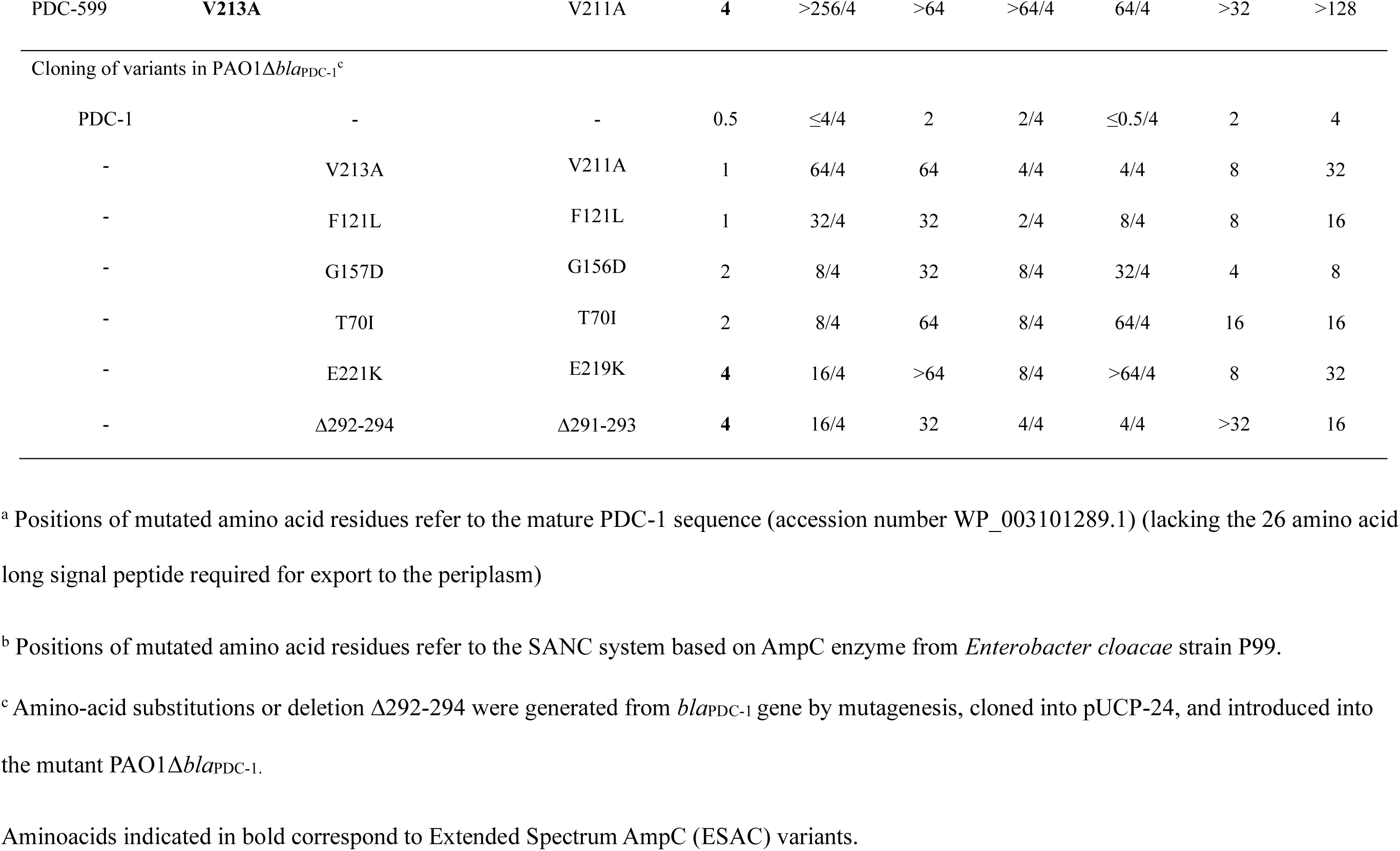
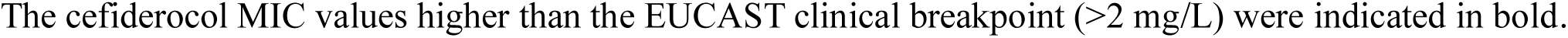
Effects of PDC variants on cefiderocol susceptibility.

### Outer membrane tonB-dependent siderophore transporters and their regulators

In addition to cefiderocol resistance by β-lactam mediated hydrolysis, various mutations that impact active tonB-dependent receptors periplasmic uptake of cefiderocol cause resistance by limiting uptake. Such mutations have been reported both in the transporter themselves (particularly PiuA (or ortholog PiuD), and PirA), or in the regulators of these transporters (8, 10). Genome analysis of the 103 cefiderocol resistance strains found 40 strains (38.8%) to have a non-synonymous nucleotide polymorphism (SNPs) in either the TonB dependent transporter or their known regulators, and these mutations were particularly frequent and often multiple in strains with high cefiderocol MIC levels (>16 mg/l) (Tables 2 and S2). Regarding TonB dependent transporters, SNP were identified in *piuA/piuD* (*n*=11, 10.7%), *pirA* (*n*=9, 8.7%), *fpvB* (*n*=9, 8.7%), *piuC* (*n*=8, 7.8%), *fecA* (*n*=8, 7.8%), *fptA* (*n*=5, 4.9%), *pfeA* (*n*=5, 4.9%) and *fiuA* (*n*=3, 2.9%), though association with cefiderocol resistance is not known in all cases. Most of these identified mutations resulted in protein truncation aberrant sequences, or complete gene deletion and thus predicated loss of transporter function. While deletions of the PiuA hydroxamate-type ferrisiderophore receptor and the PirA ferric enterobactin receptor have previously been shown to impact cefiderocol susceptibility, we further evaluated the impact of deleting *piuC*, *fptA*, *pfeA*, and *fpvA* in the PAO1 strain on cefiderocol activity (Table S3). This evaluation found that neither *fptA* nor *pfeA* played a major role in cefiderocol transport, and that loss of *fpvA* led to sensitization to cefiderocol (likely due to increased iron stress by inability to take up pyoverdin). Interestingly, deletion of *piuC*, which encodes an iron-dependent oxygenase TonB-dependent receptor, did result in significant increase in the cefiderocol MIC (from 0.125 to 2 mg/L), and may thus be a genetic factor regulating cefiderocol resistance in clinical strains. Regarding the transcriptional regulators of the TonB dependent transporter, SNPs were identified in *pirR* (*n*=16, 15.5%) and *pirS* (*n*=3, 2.9%). PirR is the response regulator belonging to the two-component system PirRS that modulates the expression of both the PirA, and PiuA (PiuD) transporter, and was the most frequently mutated transport related locus in the cefiderocol-resistant clinical strains. Interestingly, except for one strain which have a deletion of 1 nucleotide in *pirR* gene, all the strains have a frameshift mutation (R132*fs*) occurring in a region with a succession of 7 guanine residues from nucleotides 389 to 395 (amino acid position 132 or 133). The cefiderocol MICs of strains harboring the *pirR* frameshift ranged from 4 to 128 mg/L (Table 2). Of these *pirR* frameshift mutants, only one strain that exhibited a cefiderocol MIC of 8 mg/L, lacked any acquired β-lactamase or ESAC variant (ST1689, PDC-1) suggesting an important role of PirR function in clinical susceptibility to cefiderocol. Deletion of the *pirR* gene in the strain PAO1 confirmed that loss of function of PirR leads to a significant reduction in cefiderocol activity (4-fold increase in MIC) (Table S3).

## Discussion

In this nationwide study, we analyzed 103 cefiderocol-resistant *P. aeruginosa* clinical isolates collected over a four-year period, and provided a comprehensive overview of the epidemiology and mechanisms underlying resistance to this last-resort siderophore cephalosporin. Cefiderocol-resistant isolates exhibited extensive multidrug resistance, with high rates of resistance to β-lactams, carbapenems, and β-lactam/β-lactamase inhibitor combinations (ceftolozane/tazobactam, ceftazidime/avibactam, and imipenem/relebactam). This was consistent with the high proportion of strains producing carbapenemases and/or an ESBL (72.8%, *n*=75) in our collection. These findings align with recent epidemiological data showing that cefiderocol non-susceptibility is enriched among ceftolozane/tazobactam-resistant (8.3%, 95% CI 4.1-16.0%) and NDM-producing isolates (22.9%, 95% CI 8.1-49.7%) (27). In this context, colistin remained the most consistently active antibiotic, highlighting its role as a last-line therapeutic option.

Cefiderocol resistance has been reported across multiple STs, including high-risk clones such as ST111, ST175, ST235, and ST308, without any clear lineages-specific association (4, 18, 28–30). Together with our data, this supports a polyclonal emergence of resistance driven by diverse chromosomal and acquired mechanisms rather than clonal dissemination.

Although cefiderocol was initially considered stable against metallo-β-lactamases, a recent molecular study has demonstrated that NDM-1 enzyme efficiently hydrolyzes cefiderocol, whereas VIM-2 and IMP-1 exhibit limited activity (31). Several studies have reported reduced susceptibility or resistance in MBL-producing *P. aeruginosa*, particularly when additional resistance mechanisms are present (23, 24, 28, 32). Our data confirm that transferable β-lactamases play a major role in reducing cefiderocol susceptibility (Tables 2, 3, and S1). MBLs, especially NDM-1 (29.1%, *n*=30), were highly prevalent and strongly associated with elevated MICs (Table S2). Among strains with MIC values higher than 8 mg/L (*n*=41), 39.5% (*n*=17) produced NDM-1. These findings highlight the clinical importance of NDM producer strains detection, as cefiderocol use should be considered with caution in this context. Beyond carbapenemases, a diversity of class A ESBLs, particularly VEB-9, and PER-1, as well as class D ESBLs such as OXA-19, were frequently identified and contributed to reduced susceptibility to cefiderocol, as confirmed by cloning experiments. Notably, the acquired cephalosporinase PAC-1 emerged as a potent determinant, conferring high-level resistance to cefiderocol, as well as ceftolozane/tazobactam, and ceftazidime/avibactam. These results suggest that PAC-1 may represent an underappreciated contributor to resistance against siderophore cephalosporins (26).

In addition to acquired enzymes, mutations affecting the Ω-loop region of the chromosomal β-lactamase PDC, particularly E219K, have been associated with reduced susceptibility to several β-lactamams including ceftolozane/tazobactam, ceftazidime/avibactam, and cefiderocol (23, 33). We confirm the major impact of this substitution and identify additional mutations, including V211A, and a Δ291–293 deletion, further expanding the spectrum of PDC mutations associated with decreased cefiderocol susceptibility. The Δ291–293 deletion is located within the R2 binding region, which is critical for the recognition of bulky cephalosporins, and likely enhances hydrolytic activity. Although the substitution E219K remains rare among clinical strains (35 PDC-sequences of the 686 variants) its strong functional impact supports the need for molecular surveillance of PDC variants associated with cefiderocol resistance (34, 35).

Alterations in siderophore-mediated uptake pathways also emerged as a key mechanism. Loss-of-function mutations affecting TonB-dependent receptors such as PiuA/PiuD, PirA, and associated proteins have been reported in both clinical isolates and experimental models (8, 10, 30, 36–38). In our study, mutations were most frequently identified in the response regulator PirR, a finding that has been only rarely described (30, 38). A recurrent frameshift mutation R132fs in *pirR* gene was observed across multiple genetic backgrounds, suggesting convergent evolution. This particular frameshift in the *pirR* gene has also been reported in strains from a general hospital in Houston, Texas, not previously exposed to cefiderocol (30). Functional analysis confirmed that loss of PirR significantly reduces cefiderocol susceptibility, reinforcing the central role of impaired iron uptake.

High-level of resistance to cefiderocol (MIC>16mg/L) was consistently associated with the combination of at least two mechanisms involving β-lactamases together with alterations in siderophore transport systems. This highlights the synergistic interaction between enzymatic degradation and reduced antibiotic uptake.

Mutations in *ftsI*, encoding PBP-3, the primary target of cefiderocol, were more frequently observed among isolates with MIC values higher than 16 mg/L (Tables 2 and S2). However, the functional impact of the 9 different mutations identified in 28 strains (27.1%) was not explored in this study. Notably, previously characterized substitutions such as R504C and F533L, identified in 10.7%, (*n*=11) and 2 .9% (*n*=3) of our collection respectively, have been shown to increase susceptibility to cefiderocol, suggesting that target modifications may enhance antibiotic binding rather than confer resistance (39).

Finally, although cefiderocol is relatively stable to efflux, overexpression of MexAB-OprM has been associated with modest 2-fold increases in MIC (7, 22 ). In our collection, 48.5% (*n*=50) of strains carried inactivating mutations in at least one of the three major regulators of the efflux pump MexAB-OprM (*mexR*,13.7%, *n*=15; *nalC, 5.8%, n*=6; *nalD*, 33.0%, *n*=34). As *nalD* mutations are generally less frequent than alterations in *mexR* or *nalC*, their contribution is likely secondary (40). However, indirect effects on cefiderocol susceptibility cannot be excluded given the complex regulatory networks of *P. aeruginosa* (41).

Overall, our study demonstrates that cefiderocol resistance in *P. aeruginosa* is multifactorial and results from a complex interplay between acquired β-lactamases, chromosomal mutations, and alterations in iron uptake pathways. The absence of clonal clustering and the diversity of resistance mechanisms highlight the difficulty of predicting cefiderocol susceptibility in clinical practice. These findings underscore the need for continued surveillance and systematic susceptibility testing, particularly in strains producing ESBL and carbapenemases.

## Materials and methods

### Strain collection and antibiotic susceptibility testing

From the French National Reference Centre for Antibiotic Resistance (FNRC-RA) collection, we selected all *P. aeruginosa* clinical strains resistant to cefiderocol based on the current EUCAST breakpoints (MIC > 2 mg/L) (42). A total of 103 strains, isolated from 61 French hospitals between 2021 and 2024, were selected for the study. Cefiderocol MIC values were determined using broth microdilution in Iron-Depleted Cation-Adjusted Mueller-Hinton Broth (ID-CAMHB), following EUCAST recommendations, with the exception of the chelation contact time. Briefly, chelation of the cation MHB (Becton Dickinson) was conducted for 12 hours using CHELEX® resin (Bio-Rad, 10g/100mL). The resulting mixture underwent filtration and supplementation with Ca^2+^ (22.5 μg/mL), Mg^2+^ (11.3 μg/mL), and Zn^2+^ (0.7 μg/mL). Titrated cefiderocol powder (CliniSciences, Nanterre, France) was used for testing. Susceptibility testing for other antibiotics, including piperacillin/tazobactam, ceftazidime, ceftazidime/avibactam, ceftolozane/tazobactam, cefepime, aztreonam, imipenem, imipenem/relebactam, meropenem, and colistin, was performed using broth microdilution in CA-MHB (ThermoFisher Scientific) with customized Sensititre® microplates (ThermoFisher Scientific). MIC values were interpreted according to EUCAST 2025 breakpoints (42). To evaluate the impact of selected metallo-β-lactamases on cefiderocol susceptibility, the MIC of cefiderocol was determined in the presence of 100 mg/L of dipicolinic acid.

### Whole genome sequencing and bioinformatic analysis

Total DNA from cefiderocol-resistant strains was extracted using the Purelink^®^ Genomic DNA kit (ThemoFisher Scientific). DNA libraries were constructed using the Nextera XT DNA (Illumina), and sequencing of 150 bp paired-ends was performed on the Illumina NextSeq 500 sequencer (PibNet platform, Institut Pasteur, Paris). Data were processed using bcl2fastQ Conversion Software (v2.20, Illumina) to achieve an average depth of at least 80 for each nucleotide. DNA sequences (from 4 419 590 to 6 752 280 reads) were trimmed using FastP software (v0.23.2), and contig assembly and annotation were conducted using shovill-SPAdes (v3.15.4) and Prokka (v1.14.6) softwares, respectively. Sequence Type (ST) was determined using Pasteur’s MLST schema. Resistance genes were identified using the FNRC-RA pipeline, which is based on NDARO and CARD databases. Mutations in genes previously associated with alteration of cefiderocol activity or involved in iron transport or regulation (*bla*_PDC_, *piuA, piuC, piuD, pirA, pirR, pirS, fptA, fpvB, pfeA, fiuA, fecA, ftsI, mexR, nalB, and nalC*) were identified by comparing the sequences of cefiderocol-resistant strains to those of reference strains PAO1, PA14, CF39S, and a large collection of clinical strains susceptible to cefiderocol (MIC ≤ 0.5 mg/L) (*n*=485 strains) to suppress natural polymorphisms using CLC Genomic Workbench (v10.1.1) (43). Mutations found to be identical in more than two strains belonging to different STs were considered polymorphisms and were filtered out. The CF39S strain was chosen as the reference strain because the *piuD* gene (orthologous to *piuA*) is absent from the genomes of PAO1 and PA14. All genome sequences are available at the accession number SUB14891844.

### Cloning of interest genes into the pUCP24 plasmid

Total DNA from clinical strains was extracted from a bacterial colony using the PureLink Genomic DNA kit (ThermoFisher Scientific). Selected genes were amplified with specific primers and cloned into a linearized vector using the NEBuilder® HiFi DNA Assembly kit (New England Biolabs), following the manufacturer’s instructions. These were then cloned into the shuttle vector pUCP24 via restriction enzymes XbaI NEB® (for genes *bla*_OXA_, *bla*_PAC-1_, and *bla*_VEB_), or HindIII and EcoRI (for the *bla*_PDC_- genes). Mutagenesis of gene *bla*_PDC-1_ was performed using the Q5 site-directed mutagenesis Kit (New England Biolabs). Recombinant vectors were transferred into the PAO1 strain or its derived mutant PAO1Δ*bla*_PDC-1_ by electroporation (MicroPulser Electroporator, BioRad®). Transformants were selected on Mueller-Hinton agar medium supplemented with gentamicin (50 µg/mL).

### Deletion of genes in strain PAO1

Single knockout mutants in the *piuC*, *pirR, fptA, fpvA,* and *pfeA,* genes were constructed in the reference strain PAO1 using overlapping PCR and recombination events, as previously described (44). Briefly, fragments of −500 -bp flanking the target genes were amplified with specific primers. These amplicons were then used as templates for overlapping PCR with external primers to amplify the recombinant fragments. The resulting amplicons were cloned into a linearized vector using the NEBuilder® HiFi DNA Assembly kit (NEB) and subsequently subcloned into the suicide vectors pKNG101 or pFOG and *E. coli* CC118λ*pir* as BamHI-HF or BamHI-EcoRI fragments. The recombinant vectors were then transferred into the PAO1 strain by conjugation with the pRK2013 vector, and deletion mutants were selected on *Pseudomonas* Isolation Agar supplemented with 2,000 µg/mL streptomycin. Deletion mutants were then isolated on minimum M9 supplemented with 5% sucrose. Both strands of the DNA were sequenced to confirm the deletion.

## Acknowledgments

We thank all French laboratories that contributes to the surveillance of resistance in *Pseudomonas aeruginosa.* This work was funded by the French Ministry of Health though the Santé Publique France agency.

